# A Biosynthetic Thumb Prosthesis

**DOI:** 10.1101/2025.05.23.655850

**Authors:** Sachi Bansal, Gracia V. Lai, Marcus Belingheri, Amber Q. Kashay, Jinyoung Kim, Alyssa Tomkinson, Samantha Herman, Keval Bollavaram, Brian K. Zukotynski, Nirbhay S. Jain, Kodi K. Azari, Lauren E. Wessel, Tyler R. Clites

## Abstract

In cases of severe damage to the extremities, the function and structure of compromised biological tissues must be replaced. If biological reconstruction using autologous tissue is not feasible, amputation and replacement with a synthetic prosthesis is often the next best option. Currently, the synthetic materials available for prostheses are limited, especially in their ability to restore skin sensation. Here we show a biosynthetic prosthesis that combines the versatility of titanium with the rich sensory capabilities of biological skin, to both increase movement and restore sensation to an amputated thumb. The prosthesis recreates opposition pinch by linking motion of the prosthetic joint to that of the residual biological joint. The device is enclosed in neurotized skin from the patient’s own body, providing natural sensation. We validated the biosynthetic thumb’s ability to reproduce opposition pinch on the benchtop and in a cadaver, and showed *in vivo* viability of the skin interface in an animal model. These results provide a framework for functional reconstruction of amputated digits using a combination of synthetic materials and biological tissues.

**One Sentence Summary:** A prosthetic thumb made up of a titanium mechanism and living skin can restore opposition pinch and touch sensation following thumb amputation.

## INTRODUCTION

The thumb is considered clinically to be the most important digit on the hand, responsible for approximately 40% of hand function ^1–4^. The thumb’s core role is opposition, which is critical for grasp and pinch ^5^. Natural opposition requires an ability to trace the tip of the thumb along the tips of the remaining four fingers. In the intact hand, this ability is a direct result of thumb’s position on the hand and complex anatomical structure ^5^. The interphalangeal (IP) joint, which is the thumb’s most distal joint, plays a crucial role in functional opposition pinch, accounting for up to 31% of pinch strength and ensuring fingertip-to-thumb-tip contact ^6,7^. The rich sensory information from the skin at the thumb tip allows for robust closed-loop control of pinch and grasp force, as well as perception of texture, slip, and temperature ^8^. The skin also acts as a barrier to external insult such as infection.

Catastrophic injury to the thumb is fairly common: approximately 6 million digital amputations occur annually worldwide, one third of which include loss of at least one thumb joint ^9^. Because the thumb plays such a critical role in hand function, its loss can have tangible effects on lifestyle ^10^. This is especially true in populations who rely on their hands for manual labor, which incidentally predisposes them to these very amputations; loss of the thumb can inhibit their ability to perform work functions, creating financial strain or necessitating a shift in career. More broadly, partial hand amputation is linked directly to decreases in mental health and overall quality of life, with a large portion of the deficit related to functional impairment ^10^. While biological reconstruction using native tissues is typically favored, it is not always possible. In such cases, we face the task of reproducing the function of the lost biological digit via synthetic prostheses. For some of these functions (e.g. actuation, mechanical strength), synthetic components (motors, titanium) outperform the biological systems (muscles, bone) that they are intended to replace ^11–13^. However, other functions, such the skin’s ability to sense the environment and protect from infection, have proven particularly challenging to replicate with synthetic components ^14–16^. As a result, it has been difficult to create synthetic digits that look, feel, and behave like their biological counterparts ^17–19^.

With no good option for autologous reconstruction, thumb prostheses face the challenges of restoring functional opposition pinch, attaching to the hand, and providing natural touch sensation. Conventional external prostheses, such as the GripLock Finger (Naked Prosthetics, Olympia, WA, USA), are effective in restoring opposition pinch but cannot provide cutaneous feedback to the nervous system ^20,21^. This type of prosthesis typically straps to the hand, which is not a rigid attachment and adds bulk that can obstruct hand function. Percutaneous osseointegrated implants provide a low-profile option for rigidly attaching prosthetic thumbs to the residual bone; however, osseointegrated thumb prostheses do not have a functional IP joint, and therefore cannot restore opposition pinch. Without neural integration these devices also cannot restore skin sensation, and the percutaneous abutment creates a chronic infection risk ^22^. Even the most advanced of current clinical devices offer only partial restoration of function, and none have successfully replicated the critical tactile feedback and protective properties of native skin. In fact, the shortcomings of existing prosthetic technologies are significant enough that some patients choose to sacrifice another of their digits to restore thumb function (e.g. toe-to-thumb transfer). While this purely biological approach restores both opposition pinch and sensation, toe-to-thumb transfer is a technically demanding surgical procedure, provides a limited range of motion compared to the native thumb, and can lead to changes in gait due to donor site morbidity ^23^. Other patients forego prosthetic reconstruction altogether and instead adapt to life without a thumb ^24^, which comes at significant functional cost ^1–4^.

Synthetic “skin” and direct neural interfacing have shown promise in experimental devices as a means of restoring cutaneous sensation, but still cannot compete with the density, precision, or versatility of sensory end organs in biological skin ^15,25,26^. In addition to sensing, *communication* of high-density information from external sensor arrays to the nervous system remains a persistent challenge ^27^. Direct nerve stimulation has been useful in restoring touch sensation from a single sensor that localizes to a region (e.g. the whole thumb-tip), and some preliminary evidence has been provided that such stimulation can evoke natural touch perception ^28–30^. While this approach has been shown to improve task performance, direct nerve stimulation faces drawbacks including low resolution and precision, uncomfortable or unnatural sensation, and sensory adaptation ^31,32^. Current nerve stimulation approaches also require either implanted electronics or percutaneous transmission of power, which can pose challenges for long-term viability. Cutaneous sensory information has also been delivered directly to the brain via intracortical microstimulation; however, this approach has faced challenges with stability and longevity and requires invasive surgery that is difficult to justify for persons with thumb amputation ^33,34^.

As an alternative to purely-synthetic reconstruction, we propose an approach in which the prosthetic thumb is reimagined as a hybrid system that selectively leverages both biological and synthetic subsystems where each is most advantageous. Specifically, we describe a biosynthetic prosthesis that combines a titanium linkage with the patient’s own skin to both enable opposition pinch and restore skin sensation. At the core of the device is a four-bar linkage, which couples motion of the prosthetic IP joint to that of the biological metacarpophalangeal (MCP) joint; this design provides opposition pinch without the complexity of powered actuators and does not require attachment of tendon to synthetic material, which remains a significant unsolved problem ^35–37^. Although fully-implanted articulating prostheses are common in orthopaedics ^38,39^—and have even been proposed as a solution for leg amputation ^40^—their movement depends on attachment of tendon to metal or other synthetic material. In our system, the metal portion of the biosynthetic thumb anchors only to *bone*, providing robust long-term attachment and actuation. The biological portion of the prosthesis comprises the patient’s own vascularized, innervated skin, sustainably harvested from elsewhere on the arm and hand. Current endoprosthetic devices are designed to be buried beneath a robust soft tissue envelope ^38,39^, and therefore cannot be applied to treat pre-existing or traumatic amputations ^40^. In contrast, our biosynthetic thumb incorporates transposed autologous skin onto the prosthesis itself, eliminating the need for existing native skin at the site of reconstruction. This transposed skin becomes the outer surface of the prosthesis, protecting the body from infection and leveraging innate cutaneous end organs to provide natural touch and temperature sensation.

This manuscript describes comprehensive preclinical validation of the biosynthetic thumb, addressing surgical, mechanical, and clinical feasibility. We first use a cadaver to establish and validate a new surgical paradigm for transposition of the patient’s skin onto the surface of the device. We then present an optimization and evaluation of the device’s mechanics in free space and in the context of opposition-relevant loading. We describe an animal study of the proposed synthetic architecture in living skin, through which we establish long-term clinical viability of the skin-implant interface. The manuscript concludes with a surgical and functional evaluation of the complete implant in a human cadaver model.

## RESULTS

### Surgical approach to covering the prosthesis with living skin

To create a biosynthetic prosthesis with real translational potential, it was first necessary to develop a surgical approach that would allow the implanted component to be *entirely* covered in living skin. Complete coverage of the implant (in contiguity with the skin of the residual thumb) ensures a sealed skin envelope, which is crucial to keeping the transposed skin alive and preventing infection. This surgical context became the starting point for our design process; we deliberately chose to design the implant around the surgery (and not *vice versa*), to increase the likelihood that the prosthesis would be clinically viable and work synergistically with the body to restore function. Through a series of cadaver dissections, we developed and demonstrated a new surgical approach that combines established flap techniques in a new way such that they can be used to cover our fully-synthetic implant core (Fig. 1a).

**Figure 1.**
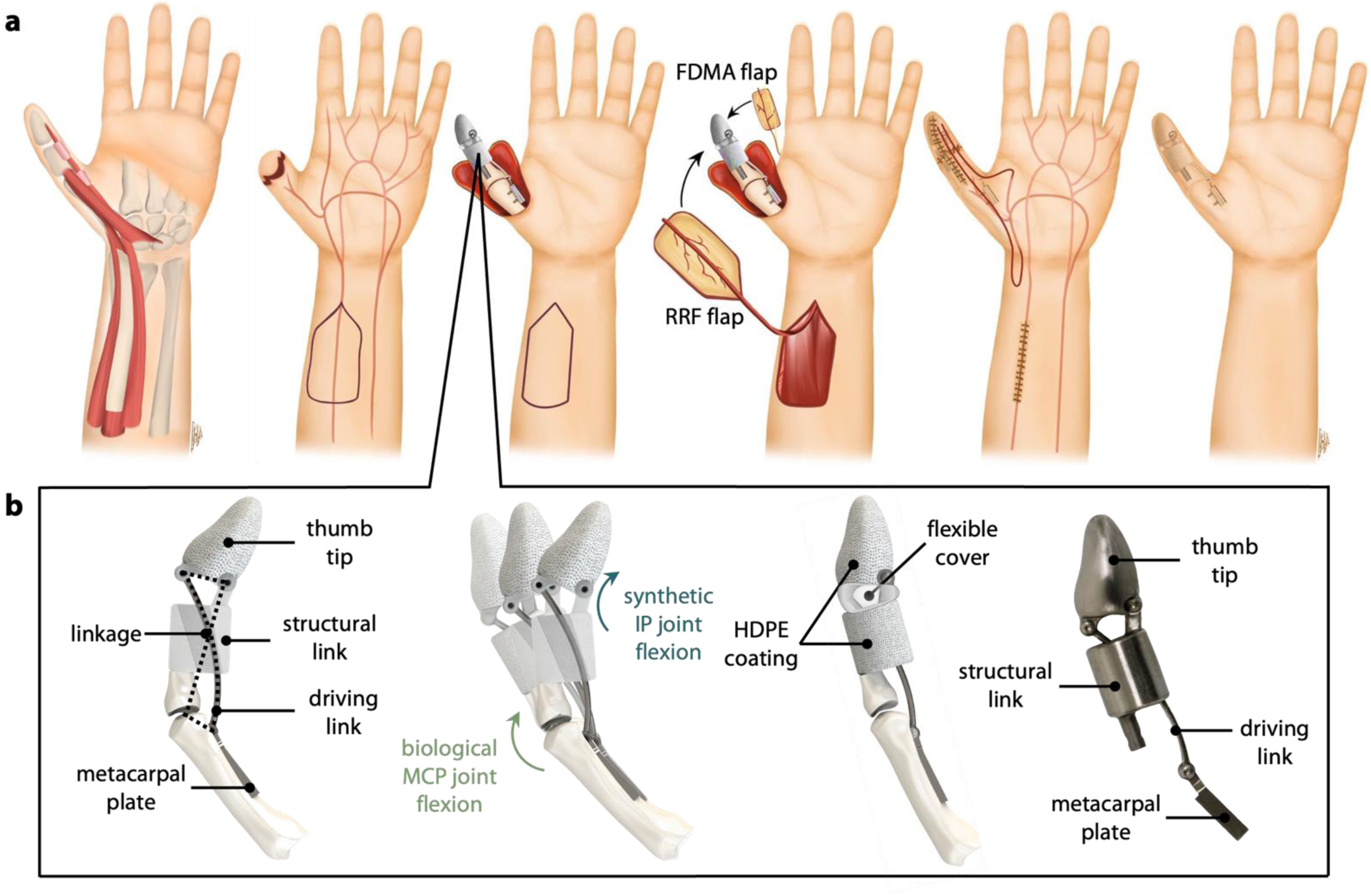
Surgical approach and device design. **a.** (left to right) Biological thumb with intact native joints and soft tissue. The IP joint is amputated, and native skin covers residual proximal phalanx. The device is attached to the local bony structures including the residual proximal phalanx and metacarpal. The reverse radial forearm (RRF) and first dorsal metacarpal artery (FDMA) flaps are harvested and rotated about their pedicles to cover the implant. The donor and recipient sites are sutured closed, and the system is allowed to heal. **b.** The implant is made up of four parts: the thumb tip, structural link, driving link, and metacarpal plate. The linkage connects the MCP and IP joints, such that flexion of the biological MCP joint creates flexion of the synthetic IP joint. Static (non-joint) portions of the implant are covered in porous HDPE, and the synthetic IP joint is protected by a flexible cover. A prototype was manufactured in Ti-6Al-4v.

The intramedullary canal of the residual proximal phalanx was first identified as the primary site for bony anchoring, with the driving link secured to the volar aspect of the metacarpal using two biocortical screws (M2) through the metacarpal plate. We selected a combination of two cutaneous flaps to cover the device with innervated, vascularized autologous skin. The first of these is the RRF flap, which provides surface area to surround the implant. A RRF flap measuring 8 cm by 4 cm was harvested from the forearm ipsilateral to the thumb amputation. The proximal end of the radial artery within the flap was clipped and ligated; in a living patient, this would induce retrograde flow in the radial artery, allowing the flap to be perfused from its distal end without compromising flow to the hand ^41^. The RRF flap on its vascular leash was then passed through a subcutaneous tunnel in the wrist, tubularized around the thumb implant, and sutured to the distal edge of the native skin surrounding the residual thumb. The portion of this RRF flap over the thumb tip was then replaced with an innervated 1.5 cm x 2 cm first-dorsal metatarsal flap (FDMA flap) from the dorsal aspect of the index finger; this flap is used to provide coverage and restore sensation to large wounds on the tip of thumb ^42,43^. A complete depiction of the surgical technique in the context of the full thumb prototype is shown in Supplemental Fig. 1.

In these dissections, both flaps were successfully harvested, rotated on their respective vascular pedicles, and reattached to the residual skin at the base of the thumb. The skin flaps were large enough to cover a proxy implant without excessive tension in the transposed skin. The defects created on the forearm and wrist were sufficiently small to be sutured closed by primary intention (Fig. 1a); the defect on the index finger from the FDMA flap is usually closed via a split-thickness graft from the forearm ^44^. Because these flaps are not typically used to cover synthetic implants, extrusion through the reconstructed skin was a significant concern ^45–49^. We addressed this by covering the non-joint portions of the device, which do not move relative to the surrounding skin, in porous high-density polyethylene (HDPE) to promote skin ingrowth during the healing process ^50,51^. In order to prevent soft tissue impingement or infiltration that could disrupt link movement and joint motion, we determined that future iterations will include flexible silicone covers over all articulations (Fig. 1b). Silicone was selected for its flexibility and history of use in orthopaedic applications (e.g. Swanson Flexible Finger Joint Implant, Stryker, Kalamazoo, MI, USA), and in pacemakers ^52^ to protect the device from the surrounding biological environment.

### Optimization and validation of mechanical linkage

The synthetic portion of the implant is a crossed four-bar linkage that couples motion of the prosthetic IP joint to that of the biological MCP joint. Importantly, this motion is enforced through bone-anchored metal components, which avoids the need for tendon-metal interfaces; such soft-tissue interfaces are notoriously prone to failure in the body ^35–37^. This linkage was optimized to match a target trajectory derived from a published relationship between MCP and IP motion in the biological thumb during opposition pinch ^5^, while accounting for geometric constraints of the anatomical envelope. An initial optimization yielded an “idealized” linkage design, with a mean absolute percent error (MAPE) of 0.73% relative to the target angle relationship (Fig. 2). However, the two links that formed the IP joint extended beyond the device envelope, posing a high extrusion risk. As such, we constrained the links to remain within the diameter of the “structural link” (Fig. 1b) and re-optimized. This constrained optimization produced the “clinically viable” linkage design, which had an MAPE of 15.2% from the ideal angle relationship. Specifically, the slope of the MCP-IP relationship was shallower, resulting in an IP joint that was 3.4° less flexed at the minimum MCP joint angle, and 4.4° more flexed at the maximum MCP angle (Fig. 2b, blue line).

**Figure 2.**
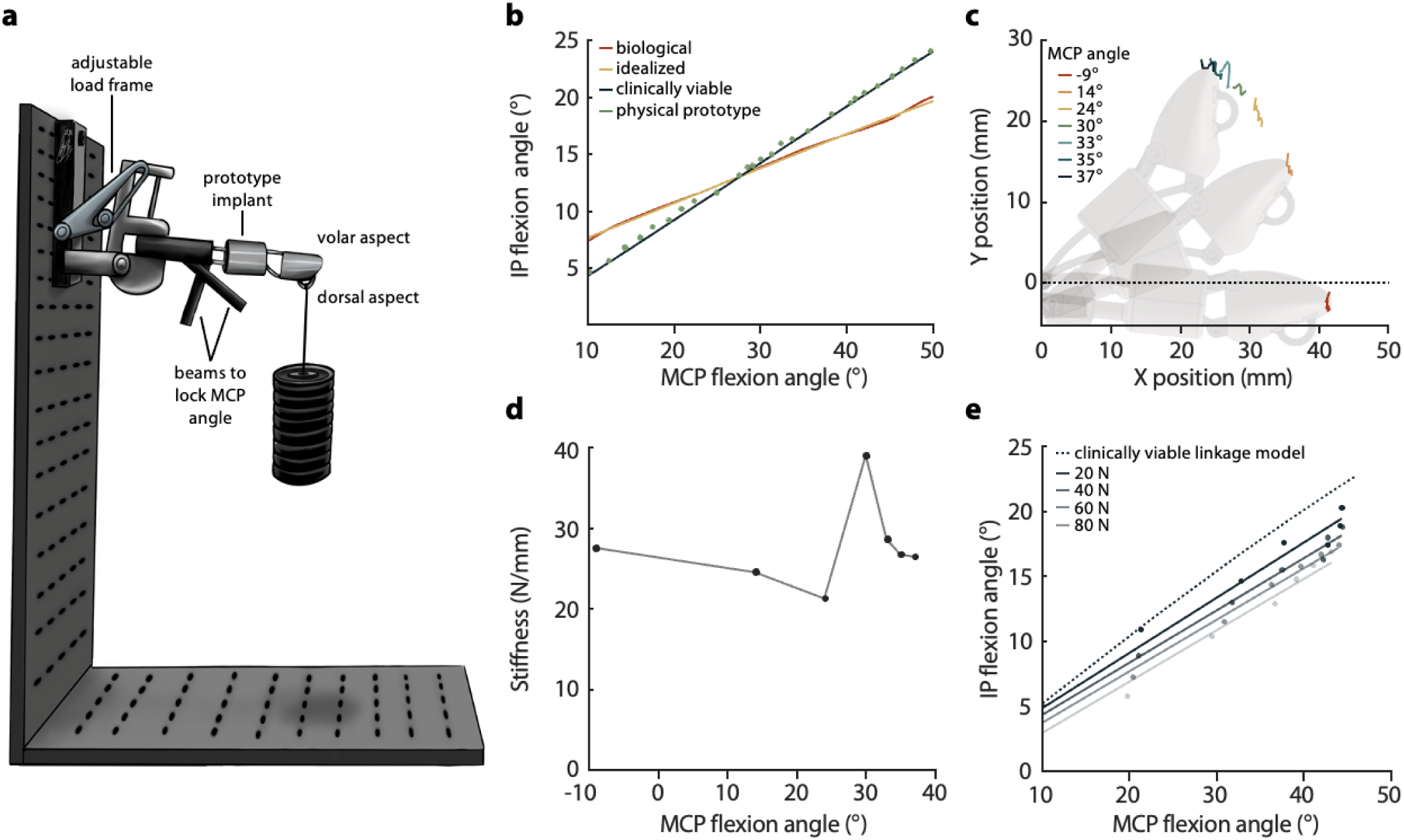
Benchtop mechanical testing results. **a.** Experimental setup, in which the prototype thumb is clamped into an adjustable load frame and loaded with the proxy MCP joint locked at different angles. **b.** MCP-IP angle relationships showing the target trajectory, the idealized linkage design, the clinically viable linkage design, and the experimental performance of the physical prototype (Supplemental Table 1.). **c.** Deflection of the thumb-tip marker across seven locked proxy MCP angles. **d.** Stiffness of the device as a function of MCP angle. Stiffness represents the amount of perpendicular force required at the thumb tip to produce 1 mm of linear deflection of the thumb tip. **e.** Experimental MCP-IP angle relationship in the presence of different loads. Dots show average measured angle over 1.5 seconds, and solid lines show first-order linear regression fits. The dashed line represents model predictions for the clinically-viable device (identical to the solid blue line from b.).

A prototype of the clinically viable linkage design was manufactured in Ti-6Al-4V, and attached at its distal end to a simple mechanical hinge (proxy MCP joint). This prototype was then affixed to an adjustable loading frame, and loads were applied to a hook on the dorsal aspect of the thumb tip (Fig. 2a). In the absence of external loads, the proxy MCP joint and the device’s IP joint were able to flex over a range of 70.3° and 29.7° from their rest positions, respectively (Fig. 2b, green dots), and matched the predicted kinematic relationship with 2.84% MAPE.

Thumb-tip loads during a typical grasp activity can exceed 80 N ^53–55^. With the linkage locked at seven different proxy MCP angles (Fig. 2c), ranging from -9.0° (slight extension) to 37.1° (flexion), we observed a maximum of 3.4 mm of deflection at the thumb tip under 82.4 N of force (Supplemental Fig. 2a). Angular stiffness about the base of the implant changed as a function of proxy MCP angle, with a maximum effective deflection of 9.4° under 82 N of force perpendicular to the thumb tip, corresponding to a moment of 3.1 Nm about the proxy MCP (Supplemental Fig. 2b). Stiffness of the device varied as a function of proxy MCP angle and was highest with the proxy MCP and IP near the middle of their ranges of motion (Fig. 2d). Deflection of the thump tip decreased the effective IP angle by up to 5.87° under 82 N, shifting the MCP-IP relationship toward less IP flexion (Fig. 2e).

### Validation of skin-implant interface in an animal model

Although the proposed surgical techniques are part of established clinical practice, these flaps are not typically used to cover synthetic implants in an area of high motion. This introduces concerns of flap viability and potential extrusion. To address these concerns, a rodent model was used to validate long-term viability of a proxy implant in the context of a large tubularized rotation flap. In a total of 23 rats, vascularized skin flaps were elevated, tubularized, and rotated to a new orientation in a high-motion area near the hind leg (Fig. 3a). In 12 rats, the flap was wrapped around a proxy implant (experimental group); the remaining 11 did not have implants (control group). Proxy implants were two-part components shaped analogously to the thumb implant and made of the same HDPE material (Supplemental Fig. 3). Two rats from each group were lost due to flap detachment caused by chewing. In the remaining 19 animals, flaps were allowed to heal for 3.5 to 11.25 months. After the sutures in each animal had dissolved, we assessed strain in the flaps as the rodents walked freely across a transparent cage (Movie S1); these trials served to ensure that the flaps deformed each time the animal walked, in a way that adequately represented the large amount of skin motion in the human thumb. All flaps exhibited large strains, with averages ranging from 69% to -17% (Supplemental Fig. 4).

**Figure 3.**
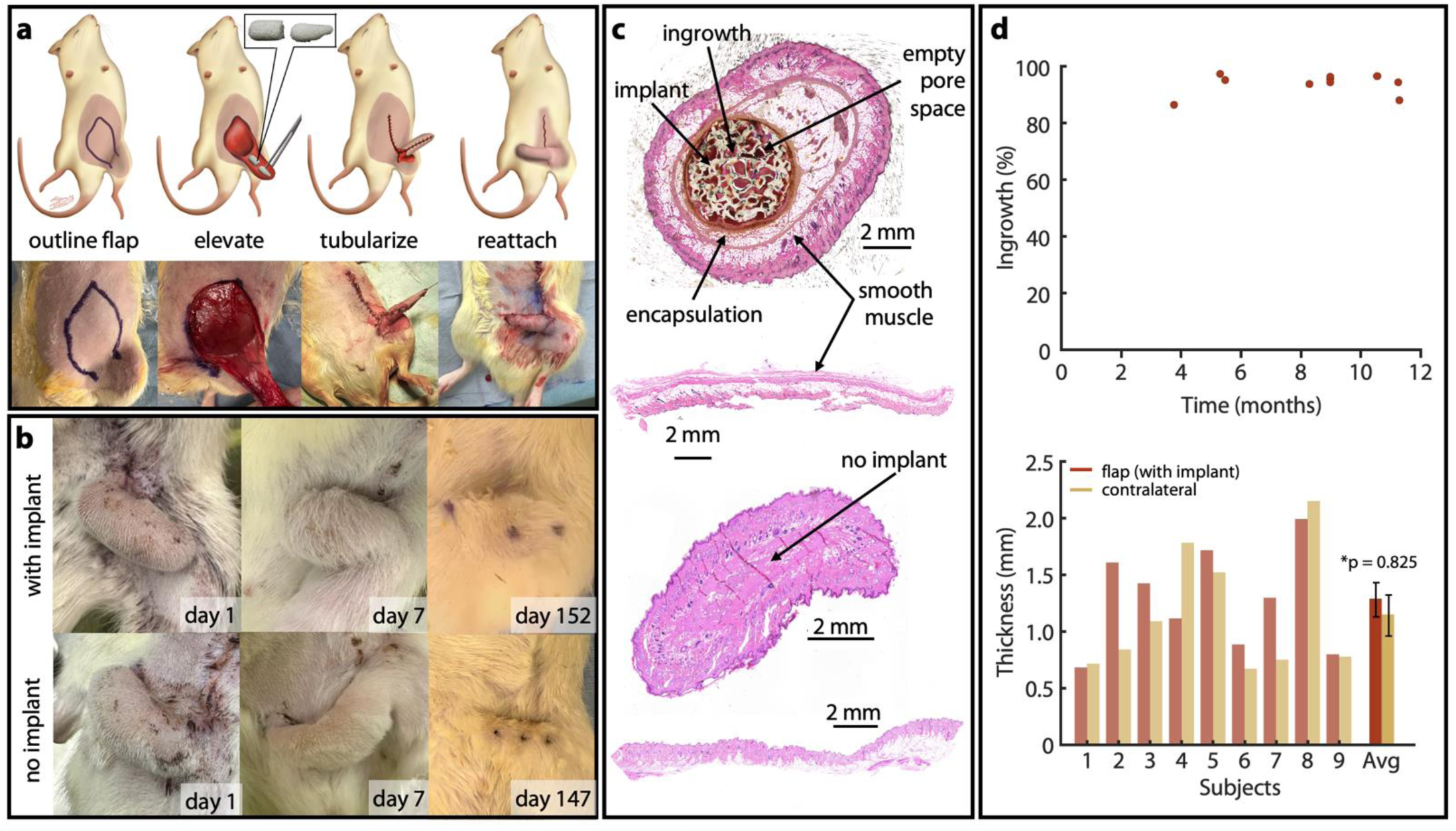
Surgical design, post-op follow-up, and histological analysis from animal study. **a.** Rodent surgical technique for both experimental (with implant) and control (no implant) groups. **b.** Post-op photos from immediate post-op through tissue collection. **c.** Representative histology. Top two samples are from the same animal (with implant), taken from the flap (top) and the contralateral epigastric region (bottom). Bottom two samples are from a second animal (no implant), again from the flap (top) and the contralateral epigastric region (bottom). **d.** (top) Percent ingrowth versus time at tissue harvest, where each point represents one animal. (bottom) Smallest distance from external surface of epidermis to external surface of smooth muscle layer for the skin flap (with implant) and contralateral skin sample.

Our primary measure of success in these experiments was lack of extrusion; according to this metric, we observed a 100% success rate in the animals with implants. All flaps in both groups healed well, and showed no gross signs of necrosis or tissue damage (Fig. 3b). Qualitative histological analysis in the experimental group revealed an implant surrounded on all sides by healthy dermal tissue, with some pockets of subcutaneous fat (Fig. 3c). By volume, tissue filled 94.3% ± 3.85% of the implant voids. We did not observe any relationship (R^2^ = 0.02) between ingrowth and post-op time prior to tissue harvesting (Fig. 3d). We found no significant difference in skin thickness (p = 0.825, Fig. 3d, right) between the experimental flaps and skin samples collected from the contralateral side of the epigastric region (Fig. 3c, bottom), suggesting that neither the presence of the implant nor tubularization of the skin flap caused thinning of the surrounding flap tissue (Fig. 3d, bottom). We also evaluated tissue encapsulation, which we defined as the proportion of the implant’s circumference covered by thick fibrous connective tissue; by this metric we observed 100% encapsulation in all implants (Supplemental Figs. 5, 6).

### Performance of the complete biosynthetic thumb in a cadaver

We used a cadaver to characterize opposition pinch after attachment of the implant to the residual proximal phalanx. In the cadaver (Fig. 4), thumb function was compared between two conditions: 1) the native biological thumb, actuated by pulling on the flexor pollicis longus (FPL) in its native insertion (Fig. 4a, left); and 2) the biosynthetic thumb, comprising the uncovered implant anchored to the residual bone and actuated by pulling on the FPL in its post-amputation, transposed insertion (Fig. 4a, middle; Supplemental Table 1). We also completed the reconstruction by covering the thumb implant with the proposed flaps (Fig. 4a, right); however, we did not include this condition in our biomechanical experiments because as confirmed radiographically, motion of the implant *within* the skin envelope produced an inaccurate representation of IP motion. We do not expect that this relative motion will affect performance of the thumb *in vivo*, due to ingrowth of the skin into the porous HDPE coating. In the cadaver, the two flaps provided sufficient coverage to completely encapsulate the implant, and the reverse radial forearm (RRF) flap donor site was small enough to be closed via primary intention (Fig. 4a, right). When pulling only on the FPL tendon and finger flexor tendons, the fully-reconstructed thumb created flexion movement at both the biological MCP and synthetic IP joints (Fig. 4b, Movies S2 and S3) that was sufficient for functional opposition pinch (Fig. 5).

**Figure 4.**
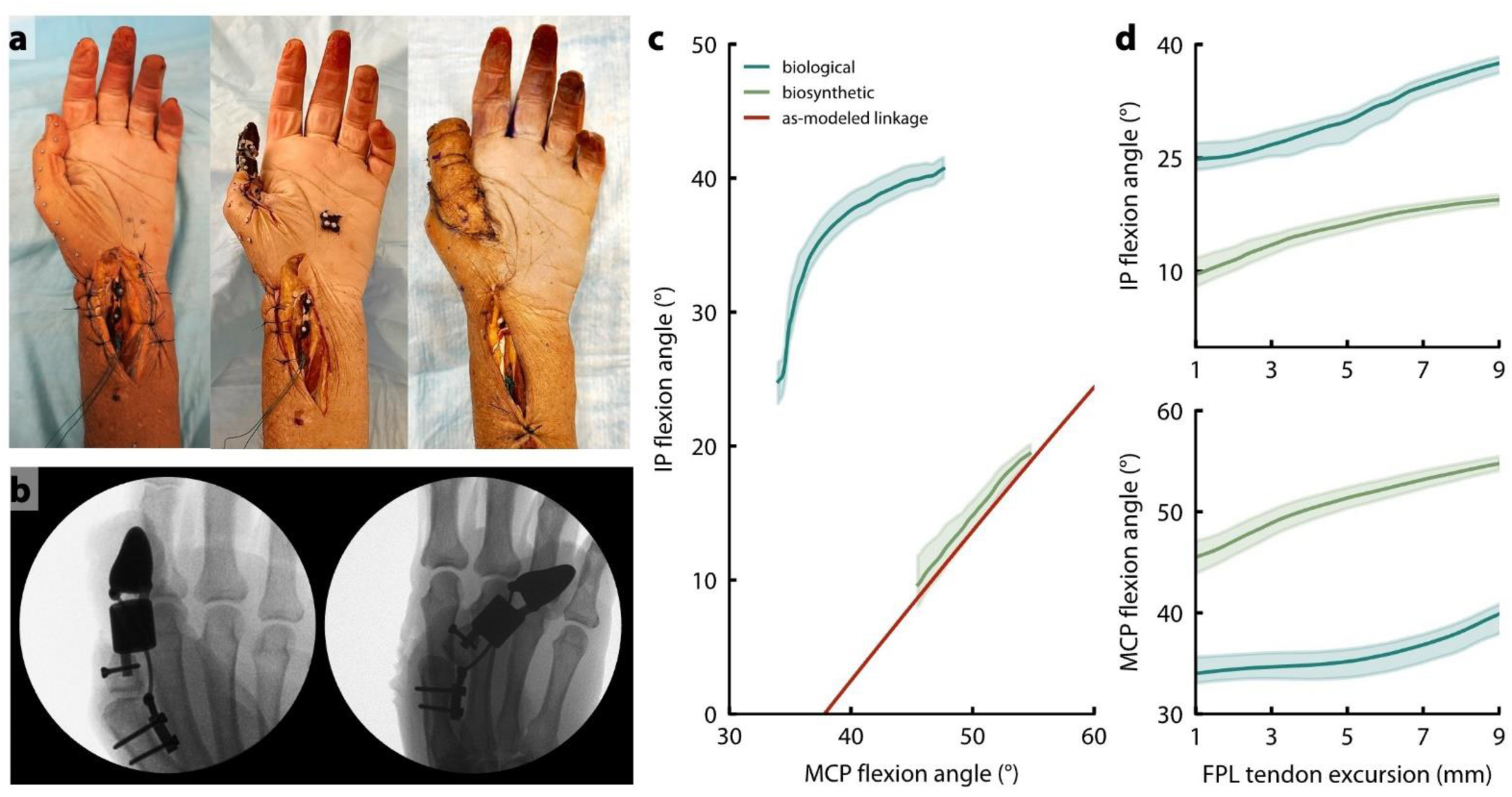
Cadaver testing results. a. Cadaver hand with biological thumb (left), biosynthetic thumb (middle), and biosynthetic thumb with flap coverage (right). b. Radiographic images showing the biosynthetic thumb with flap coverage in the rest position (left) and at full flexion (right). c. MCP-IP angle relationship for the cadaver hand with biological thumb and with biosynthetic thumb, when actuated only by pulling on the FPL tendon. Solid blue and green lines represent the mean of 10 trials, and shaded regions show +/- 1 standard deviation. Predictions from the as-implanted linkage model are also shown for comparison (red). d. IP (top) and MCP (bottom) flexion angle as a function of FPL tendon excursion. Zero excursion represents resting tension in the FPL tendon. Solid lines represent the mean of 10 trials, and shaded regions show +/- 1 standard deviation.

**Figure 5.**
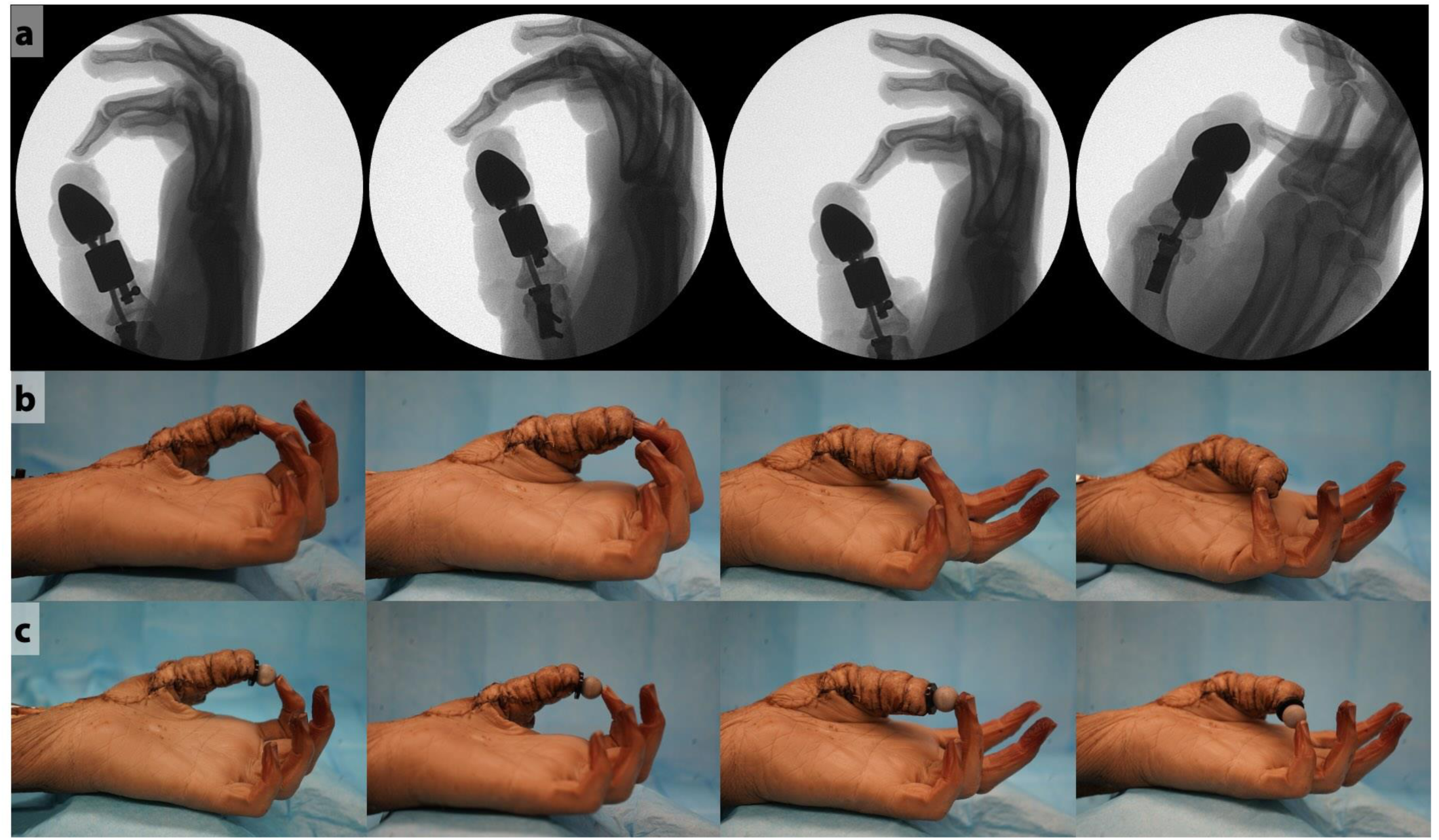
Opposition pinch with biosynthetic thumb, after coverage by FDMA and RRF flaps. **a.** Radiographs showing opposition pinch between the biosynthetic thumb and digits 2-5. **b.** Photos showing opposition pinch between the thumb and digits 2-5 (left to right). **c.** Pinch grasp holding a small spherical object between the biosynthetic thumb and digits 2-5 (left to right).

When actuated by manually excursing the FPL tendon, the IP joint in the biosynthetic thumb moved 1.08° per 1° of MCP joint flexion (Fig. 4c, green), compared to 1.12° per 1° of MCP flexion in the cadaver’s biological thumb (Fig. 4c, blue). Across a range of 9 mm of FPL excursion, the MCP and IP in the biosynthetic thumb produced 72.9% and 77.2% respectively of the range of motion observed in the biological thumb. The cadaver’s biological MCP moved between 34.0° and 47.6° of flexion (13.6° range), and the IP moved between 24.8° and 40.7° of flexion (15.9° range). Meanwhile, across about 9 mm of tendon excursion, the biosynthetic MCP moved between 45.5° and 54.8° of flexion (9.3° range), and the IP moved between 9.7° and 19.5° of flexion (9.8° range).

Although the slope of the MCP-IP joint relationship was preserved in the reconstruction, the biosynthetic IP was around 15° less flexed at baseline than the biological IP. Through post-hoc radiographic analysis of the as-implanted linkage geometry, we attributed this shift to a 6 mm difference in the distance from the MCP joint center to the metacarpal plate, between the optimized clinically viable linkage and cadaver implementation (Supplemental Fig. 7). Updating the linkage model to reflect this change resulted in agreement between the model prediction (Fig. 4c, red) and the empirical kinematics (Fig. 4c, green). Per 1 mm of FPL tendon excursion, we observed 1.05° MCP flexion and 1.22° IP flexion in the biological system and 1.03° MCP flexion and 1.09° IP flexion in the biosynthetic system (Fig. 4d). Correlation in joint flexion angle as a function of tendon excursion between the biological and biosynthetic systems was high for both the MCP (r = 0.89, p <0.0001) and IP (r = 0.99, p < 0.0001) joints, indicating a high degree of similarity between pre- and post-operative joint kinematics.

## DISCUSSION

In this study, we demonstrated feasibility of a hybrid prosthetic architecture that combines synthetic components and biological tissues to restore function, sensation, and the protective qualities of skin in a reconstructed thumb. Our new surgical approach allows for complete coverage of the thumb in vascularized skin, with natively-innervated finger skin on the thumb tip. The synthetic linkage is capable of reproducing the joint kinematics needed for opposition pinch, both in free space and in the context of the typical thumb loads. An analogous implant did not extrude or erode the skin in a high-motion area of an analogous animal model. The prototype device supported opposition pinch in a cadaver when the MCP was actuated by the native tendon.

The surgical approach by which the device is covered in native skin was designed as a new combination of established techniques. Both the RRF and FDMA flaps are commonly used to address hand, and specifically thumb, defects. ^56–58^. Our results showed that sufficiently large flaps could be harvested from these two sites to fully cover the device. Cutaneous sensation from the reconstructed thumb tip is provided through the neurotized FDMA flap, which naturally projects to the dorsal index finger ^42^. Utility of this flap in restoring natural thumb sensation has been established over decades of clinical experience ^42^. We note that sensation from the natively- innervated FDMA flap will project initially to the back of the index finger, and that the degree of sensation may differ from the native, biological finger pad tissue; however, despite these differences from the native milieu, patients show a high degree of sensation after reconstruction with an FDMA flap ^42^. To further enhance sensation, the RRF flap used for additional coverage could be harvested along with the antebrachial cutaneous nerve branch which could then be innervated at the recipient site by coaptation to sensory branches of the distal median nerve ^59^; this would increase surgical complexity, but may provide more natural sensation across the thumb. The native skin covering is also expected to protect the patient from infection, which is a major concern for osseointegrated devices that breach the skin and create a permanent open wound.

Flexion of the MCP joint caused flexion of the IP joint in both our benchtop and cadaver studies. Although it was possible to optimize the linkage mechanism such that its motion very closely replicated the biological MCP-IP joint relationships, it was necessary to deviate from this ideal to protect the MCP joint capsule and reduce the chances of extrusion. On the benchtop, the relationship between the proxy MCP and synthetic IP joints showed close agreement with model predictions. In the cadaver, we observed that the *slope* of the relationship between the biological MCP and synthetic IP joints of the biosynthetic thumb closely matched that of the biological thumb, but that the biosynthetic IP joint angle was offset in the extension direction compared to the biological thumb. This offset was caused by sensitivity of the linkage kinematics to changes in fixation geometry (Supplemental Fig. 7). To correct this, it may be possible to design patient-specific surgical guides that ensure the device is affixed to the correct locations on the bone, or to redesign the thumb tip based on each patient’s bony anatomy (*θ_t_* in Supplemental Fig. 7). Despite deviations from the biological MCP-IP relationship, the reconstructed thumb was still able to create functional opposition pinch in a cadaver when actuated only by pulling on the native tendons (Fig. 5).

The synthetic component of the reconstructed thumb maintained functional opposition pinch in the presence of high simulated pinch forces. Most importantly, under increasing loads, the MCP- IP relationship maintained the same slope, but shifted toward less IP flexion; however, this shift was relatively small compared to the overall angular deflection of the IP joint. Interestingly, we observed that non-linearities in the linkage mechanics caused a jump in the mechanism’s resistance to deformation when the synthetic MCP joint was held at 30°; we suspect that this was caused by local changes in stiffness as a function of the orientation of the loading vector relative to the curved portion of the driving link.

The miniaturized implant did not extrude in any rodents, providing evidence as to the *in vivo* viability of our surgical approach and material selection. Histological analysis revealed that the components remained securely anchored within the flap, aided by encapsulation and tissue ingrowth into the component’s pores. We did not see any significant thinning of the skin surrounding the implant despite large strain in the flap during ambulation, suggesting that an implant made from similar material will remain stable within a skin flap, even in the context of large motions. Care was taken to ensure that the miniaturized implant would be a suitable proxy for the human implant; although the miniaturized implant did not have the metal components of the full-scale device, its shape and size compared to the thickness of the skin envelope were both more conducive to extrusion and erosion than the intended application. Note that we intentionally did not evaluate cutaneous innervation in the post-operative rodent model; because the FDMA flap is a natively-innervated rotation flap (as is the analogous flap in the rodent), post-healing sensitivity in that skin is assumed based on decades of experience with similar flap reconstruction^42,43^.

We performed the full biosynthetic reconstruction in a cadaver limb, demonstrating the ability of the reconstructed thumb to support functional thumb opposition pinch. Although the MCP-IP relationships of the device *in situ* deviated from our observations during controlled benchtop testing, our results showed meaningful IP joint flexion when actuated via the native tendon. Importantly, this motion was created solely through relative movement of bone-anchored components, bypassing the need for tendon-to-metal interfaces ^60^. We grossly observed reduced motion of the IP joint after covering the implant with the intended flaps, due to motion of the implant within the skin; this further underscores the importance of skin ingrowth into the planned porous coating on the thumb tip and structural link. We attribute increased FPL tendon excursion in the biosynthetic thumb to structural changes associated with reattachment of the FPL to the base of the proximal phalanx, which is a common step in surgical thumb amputation ^61^. This increased FPL-excursion-per-unit-IP-flexion also implies a slightly increased mechanical advantage between the FPL and the IP joint in the biosynthetic thumb as compared to the native biological architecture; we expect that this mechanical advantage will help to preserve pinch and grasp strength after reconstruction, especially if there are any neuromuscular deficits or atrophy in either the native MCP flexor (*flexor pollicis brevis*) or the FPL. We also designed the surgery to preserve both the native MCP and IP flexors, to maintain as much pinch strength as possible in the reconstructed thumb; if either of these muscles is compromised either as a result of the initial injury or treatment history, pinch strength in the biosynthetic thumb may be reduced.

This work serves as an initial preclinical validation of biosynthetic thumb reconstruction, and several changes to the device are necessary prior to clinical implementation. Specific design considerations not included in the simplified prototype include polyethylene bearings to prevent metal-on-metal wear, as well as silicone covers to protect the skin from the titanium linkage components. Additionally, performance of the complete device should be assessed during cyclic loading under a representative range of loading conditions. Future research should also explore the biosynthetic prosthetic thumb as an index procedure, rather than as a secondary surgery; this could provide an immediate solution for complex thumb injury in cases where replantation is not possible.

A single treatment that both improves mechanical function and restores natural cutaneous sensation has long remained an elusive ideal in limb and digit reconstruction. By co-engineering body and machine our approach uniquely combines the strength, machinability, and durability of synthetic material with the unparalleled sensing and protective abilities of biological skin ^62^. While this initial study is focused on the thumb, due to its critical importance to hand function, the approach could be readily adapted for reconstruction of other digits. Similar techniques are already used selectively in phalloplasty, which involves construction of a cylindrical appendage from a tubularized skin flap and often incorporates implantation of a synthetic penile prosthesis ^63–65^. These procedures have relatively low complication rates, providing additional confidence in the ability of pedicled skin flaps to heal around a synthetic implant ^65,66^. Expansion of this biosynthetic approach to larger reconstructions, such as full limb joints, would be limited by actuation and flap viability ^40^; however, these challenges could conceivably be overcome with passive joint structures or implanted actuators, and with the help of tissue-expanders, multi-stage flap pre-conditioning, and prelamination ^67–69^. By bridging the gap between synthetic and biological reconstruction, this work lays the foundation for a new class of surgically-invasive prosthetic technologies that seamlessly integrate with the body to provide natural touch sensation. If we expand our “menu” of reconstructive materials to include not only titanium struts and copper wires, but also skin and nerves, we may be able to combine the best of the biological and synthetic worlds in maximizing functional restoration.

## MATERIALS AND METHODS

### Study Design

This study was designed to experimentally demonstrate feasibility of a biosynthetic prosthesis that combines synthetic materials with biological tissues to restore function and sensation after thumb amputation. As such, the experiments explicitly address the three most critical elements of the device’s design: surgical approach to implant placement and coverage, mechanical performance and robustness to load, and *in vivo* viability of the skin-implant interface.

### Surgical Approach

All cadaver work was carried out in fresh-frozen cadavers with approval from the University of California’s Donated Body Program. Specimens were stored frozen and were thawed at room temperature for 48 hours before use. At the end of each experimental day, a saline solution was applied to maintain tissue moisture. Two cadaveric dissections were conducted to identify an appropriate surgical approach to implanting the device and covering it with native biological skin (Supplemental Fig. 1). Each dissection began with a simulated amputation of the IP joint according to current clinical practice, performed by the surgeon co-authors. Several options for fixation and coverage were then explored, using 3D printed dummy implants of the approximate size and shape of the intended device. Each potential surgical approach was assessed qualitatively by the surgeon co-authors for viability and estimated likelihood of success. This design process continued iteratively and collaboratively until a suitable surgical approach was identified. Critical dimensions of the thumb were also measured during these dissections, including size of the intramedullary canal in the residual phalanx, length of the relevant thumb segments, and distance between the MCP joint center and the intended mounting location on the palmar surface of the metacarpal bone.

### Mechanical Performance

Our evaluation of mechanical performance was intended to show similarity between implant kinematics and motion of the biological thumb during opposition pinch, and ability of the implant to maintain these relationships in the presence of large grasping forces. The prosthetic device was designed as an implanted four-bar crossed linkage, in which movement of one link (in this case, the residual proximal phalanx) generates motion in the remaining links. This configuration allows movement of the biological MCP joint to be transmitted to the distal, synthetic IP joint without requiring attachment of the residual tendon to the synthetic implant material ^70,71^.

#### Linkage Optimization

We established a target kinematic relationship between the biological MCP and synthetic IP angles, to mimic the relationship between these two joints in the biological thumb during opposition pinch^5^. We note that published data regarding MCP-IP angle relationships during grasping tasks are relatively sparse. As such, we present this target kinematic relationship not as a codified optimization target, but as a starting point to demonstrate that the linkage can be tuned to produce kinematics in the ballpark of biological joint function. The linkage was parameterized into seven parameters that fully describe its mechanics (four linkage lengths and one angle) and orientation with respect to the residual thumb (two angles). These parameters were optimized using a gradient descent approach (fmincon, MATLAB 2022, MathWorks, Natick, MA, USA) with an objective function that minimized the root mean square deviation from the target trajectory of the linkage’s MCP-IP relationship. The optimization was constrained based on the anatomy, as measured in the initial cadaver dissections. Specifically, the distance between the MCP joint center and the proximal end of the metacarpal mounting place was constrained to be at least 4.5 cm to prevent disruption of the MCP joint capsule. The distance between the two screws that connected the device’s thumb tip was constrained to ensure the device links did not protrude beyond the envelope of the device which could increase the risk of extrusion of the device over time. Linkage lengths of the optimal “clinically-viable device” were: 5.04 mm, 10.05 mm, 33.51 mm, and 34.66 mm, with an initial IP angle of 7.4°. This optimized linkage architecture was implemented into a complete implant design in Solidworks (Dassault Systems, 2022). A prototype of the device was then manufactured at custom machine shop in implant-grade titanium (Ti-6Al-4V), which was selected for its proven biocompatibility and excellent fatigue strength.

#### Implant Kinematics

Prior to mechanical testing, the prototype implant was affixed at its proximal end to a titanium hinge, representing the biological MCP joint and residual proximal phalanx. The relationship between the MCP angle (synthetic only for testing, biological in the intended use) and the IP joint (synthetic part of the implant) was characterized using a 3D motion capture system (Vantage, Vicon, Yarnton, Oxfordshire), which were positioned to track the positional changes of reflective markers affixed to the device as it was passively cycled through its full range of motion. To minimize reflection from the metal device, exposed titanium was covered with non-reflective tape. Three reflective markers were placed on each segment of the prototype, along with one at the thumb tip and on each screw. For this analysis, the MCP angle was defined as the angle formed between link C and a vertical line, while the IP angle was defined as the angle formed between the line connecting the midpoints of links A and B and the line extending from the midpoint of link B to the thumb tip.

#### Mechanical Performance Under Load

Deformation of the thumb tip was characterized under increasing loads, comparable to those experienced by a biological thumb ^72,73^. A modified version of the device was employed for load testing, which incorporated two beams that enabled rigid locking of the synthetic MCP joint (and thus IP joint) at a fixed angle using a dowel pin. The prototype was mounted to a test rig that allowed for precise orientation of the thumb tip in global space, such that weights could be hung from the thumb tip at different MCP-IP angles and hang perpendicular to the thumb-tip segment. To replicate the normal forces encountered by the thumb tip during grasping and pinching activities, the device was positioned in the test rig so that its dorsal aspect faced the ground, with weights suspended from a hook on the dorsal side. Weights were added in approximately 10 N increments until a total load of 82 N was achieved, after which the weights were gradually removed until 0 N was reached. The same motion capture setup was used to quantify deformation under each subsequent load. The experiment was performed quasi-statically: each time a weight was added or removed, the system was allowed to settle for 10 seconds before applying the next load. This process was repeated for each of the seven fixed MCP positions. Marker trajectories were analyzed to determine planar translational and rotational deformation under each load. Marker coordinates were low-pass filtered at 15 Hz, projected onto the plane orthogonal to the device’s axes of rotation, and then used to determine translation and rotation of the thumb tip, as well as effective rotation of the MCP and IP angles, under load. For this analysis, MCP angle was defined as the angle between the vectors aligned with the principal axes of the metacarpal and proximal phalanx segments, while the IP angle was defined as the angle between the proximal phalanx and the central axis of the thump-tip segment.

### In-vivo Viability

All animal work was carried out in skeletally mature, male Lewis rats (250-300g) with approval from UCLA’s Animal Research Committee. Our objective in these rodent experiments was to establish viability of the proposed skin-implant interface in the setting of a rotation flap in a high-motion area. Our primary measure of success was non-extrusion of a miniaturized, analogous implant across all animals at all experimental time points. We also quantified skin ingrowth into the porous implant material and fibrous encapsulation of the gross implant.

#### Animal Implant

A two-part miniature implant was designed to capture key features of the human-scale device, including approximate size relative to the skin thickness and radii of curvature. The miniature implant was 3D printed from porous high-density polyethylene (Anatomics, Melbourne, Australia), which is the same material we proposed as a covering for the portions of the human-scale implant that should not move relative to the skin. Grossly, the miniature implant components were designed to mimic the shape of the artificial proximal phalanx and the thumb tip. The thumb tip component had a sharper radius of curvature than the analogous component of the human implant, and the ratio of skin thickness to implant size was more aggressive in the animal model, to ensure that the animal experiment posed a higher risk of extrusion than the intended human application (Supplemental Fig. 3) ^74^.

#### Animal Surgery

Twenty-three rats underwent an experimental surgical procedure in which a racket-shaped incision measuring 2 cm x 4 cm was created in the epigastric region near the hind limb. Each flap was elevated and the blood vessel supplying the flap was identified. In the experimental group (11 rats), the two-piece implant was placed on the inferior aspect of the flap, and the flap was tubularized around the implant. In the control group (12 rats), the flap was tubularized closed without an implant. The flap was then sutured closed, rotated on its pedicle, and its proximal end was affixed to the epigastric region toward the midline. Care was taken to position the flap relative to the legs so that it would move when the rat walked. The defect resulting from the elevation of the flap was sutured closed by primary intention. The rats were monitored closely in the post-operative period and treated with NSAIDs and antibiotics. Four animals (two from each group) were lost to chewing. One attempt was made in each animal to reattach the flap; all such attempts were unsuccessful, and all four animals were sacrificed prematurely. In the remaining 19 animals, after the flaps healed, we validated experimentally that the epigastric flap moved enough to simulate motion of the skin covering the human thumb. In each animal, three marks were made along the flap’s long axis with a skin marker (Supplemental Fig. 4a). Motion of these marks was then recorded through the floor of a transparent enclosure as the rats walked freely. Relative strain between each pair of marks was then extracted (Supplemental Fig. 4b-c) via a video analysis software (Tracker Video Analysis and Modeling Tool, Open Source Physics). This strain analysis was not possible as described in two of the control rats, because the flaps had become too “stretched out” from months of ambulation to support the video analysis; in these two cases, it was assumed that the skin underwent meaningful deformation during healing, as this would have been necessary for the flap to reach that stretched-out state.

#### Gross Pathology and Histological Analysis

At the conclusion of the study, animals were euthanized to facilitate harvesting of the skin flap for gross pathology and histological analysis. In the control group, samples were collected from 3 subjects at 5.25 months post-op, 2 subjects at 6.5 months, 2 subjects at 11.3 months, and 2 subjects at 19.4 months post-op. In the experimental group, 3 subjects were sacrificed between 3 and 6 months post-op (ranging from 3.7 to 5.4 months), 3 were sacrificed at 8.9 months, 1 at 10.5 months, 1 at 10.8 months, and 2 at 11.2 months post-op. Immediately after euthanasia, the flaps were thoroughly examined for signs of extrusion. The entire flap was then excised from the animal for histological analysis. One tissue sample from the experimental group was compromised while first developing the histological processing protocol; this sample was excluded from all histological analysis, leaving 9 experimental samples (Fig. 3, Supplemental Fig. 5) Tissue samples were also collected from the contralateral epigastric region for use as a baseline for skin thickness. All samples were fixed with formalin for two days and then dehydrated using ethanol (70%). Each sample was then embedded in paraffin wax and stained with hematoxylin and eosin (Scientific Solutions, Fridley, MN, USA). The slides were digitized using a Keyence-BZ X800 microscope, and analyzed qualitatively for signs of extrusion (Supplemental Fig. 5). We then used a custom software package to segment the histology images into empty space, tissue-filled space, and implant area (Supplemental Fig. 6). All software-based segmentation was checked for accuracy by a human observer. Percent in-growth was calculated as the percentage of non-implant space filled by tissue. Skin thickness was also measured manually from the digital slide by three different human observers, skilled in microscopy, whose measurements were then averaged to provide the final thickness measurement for each specimen.

### Full System Validation in Cadaver

To prevent movement during testing, the proximal ends of the cadaveric radius and ulna were screwed to a wooden board, which was then clamped to the surface of the dissection table. The cadaver specimen was placed within the capture volume of the same motion capture system used for the mechanical testing experiments. The flexor pollicis longus (FPL) tendon of the biological thumb was identified and tagged with a suture; this tendon attaches to the base of the distal phalanx and facilitates flexion of the IP and, secondarily, the MCP joints. Reflective markers were placed on bony landmarks corresponding to the IP, MCP, and CMC joint centers, as well as on the volar aspect of the distal phalanx, proximal phalanx, and metacarpal of the thumb.

We first established a baseline for motion of the MCP and IP joints in the biological thumb. Specifically, we tracked motion of the biological proximal and distal phalanx while pulling the FPL tendon along its natural direction of action, inducing flexion at the IP, MCP, and CMC joints. We then amputated the left thumb proximal to the IP joint. During the amputation, the FPL tendon was reattached to the base of the residual proximal phalanx according to standard clinical procedures; this maintains the FPL’s contribution to flexion of the MCP joint and, through the linkage, of the reconstructed IP joint. The post of the implant was placed into the intramedullary canal of the residual proximal phalanx and an interlock screw (M2) was passed volar to dorsal through the intramedullary post with bicortical fixation. The proximal end of the linkage mechanism was fixed to the metacarpal through the plate attachment with two bicortical screws (M2). Prior to coverage with skin flaps, markers were affixed to the surface of the bare implant, and motion was again recorded while pulling on the FPL tendon. Skin flaps from the forearm and index finger were then harvested according to the proposed surgical technique, and moved to cover the device. New markers were applied to both the biological and artificial joints, as well as to the previously identified bony landmarks. Ten trials of IP joint motion were collected using the cadaver’s native anatomy, followed by ten trials with the implanted prosthetic thumb. The mean joint angle relationships and +/-1 standard deviation are shown in Fig. 4c-d. Motion capture data were low-pass filtered at 6 Hz, and MCP and IP joint angle trajectories, as well as tendon excursion, were calculated directly from marker positions.

### Statistical Analysis

Geometric optimization of the linkage was performed using gradient descent, with an objective function that minimized the root mean square deviation from the target MCP-IP relationship. Stiffness of the prototype device under load was calculated as the slope of a linear regression between deformation and load at each load magnitude. To account for slop in the mechanism, the regression was allowed a non-zero y-intercept; displacement data were then baseline shifted such that the regression line passed through (0, 0). Skin ingrowth was regressed against post-op time at harvest using a linear fit with an unconstrained y-intercept. Skin thickness was compared between the flap and contralateral side using a paired *t* test at a significance level of α = 0.05. In the cadaver study, similarity between the average trajectories of the biological and biosynthetic systems was calculated using a Pearson correlation analysis.

### List of Supplementary Materials

Fig. S1: Operative technique shown in cadaver dissection.

Fig. S2: Thumb tip deflection under loading.

Fig. S3: Two-part implant design for analogous animal model.

Fig. S4: Rodent flap strain during free walking.

Fig. S5: Histology of the skin flaps from the 9 rat subjects with implants (experimental group).

Fig. S6: Histology image processing results.

Fig. S7: Sensitivity of crossed four-bar linkage kinematics to design and implantation parameters.

Movie S1: Rodent flap strain while freely walking.

Movie S2: Biosynthetic thumb motion while excursing FPL tendon (HD video)

Movie S3: Biosynthetic thumb motion while excursing FPL tendon (fluoroscopy)

Table. S1: Descriptions of all experimental groups across all studies.

## Acknowledgements

We extend our heartfelt gratitude to Dr. Joanne Zahorsky-Reeves and the UCLA Division of Laboratory Animal Medicine (DLAM) team for their support and guidance during the animal studies. We would like to thank Dr. Karen Lyon’s lab at UCLA for their assistance in the initial stages of histological processing. We extend a special thanks to Alyssa Tomkinson for her exceptional design of the illustrations featured in the figures of this paper. Finally, we wish to thank individuals who donated their bodies and tissues for the advancement of education and research.

## Author Contributions

Conceptualization: TRC, LEW, KKA

Methodology: TRC, LEW, KKA, GVL, MB, SB

Investigation: SB, LEW, KKA, AQK, JK, MB, NSJ, GVL, BKZ

Data Processing: SB, KB, AQK, TRC, SH, BKZ

Visualization: SB, AT, TRC

Funding acquisition: TRC

Writing – original draft: SB, TRC

Writing – review and editing: SB, MB, AQK, JK, AT, SH, KB, BKZ, NSJ, KKA, LEW, TRC

All authors approved of the final version of the paper.

## Data and Materials Availability

All data reported in the paper are included in the manuscript or available in the Supplementary Materials.

