## Supplemental Figures and Table for "A Biosynthetic Thumb Prosthesis"

### SUPPLEMENTARY MATERIALS

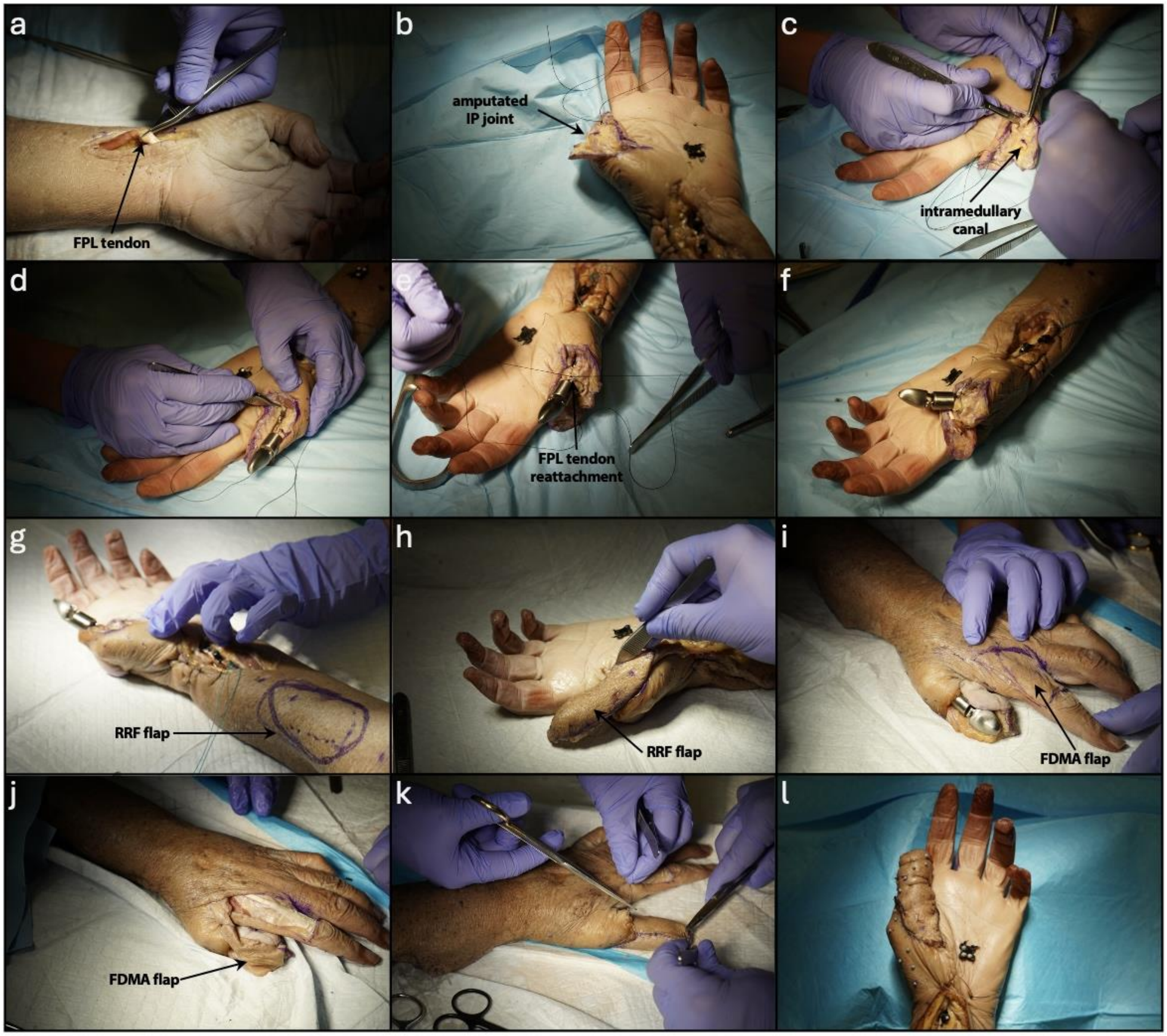


**Supplemental Figure 1.** Operative technique shown in cadaver dissection. **a.** The flexor pollicis longus (FPL) tendon is exposed on the volar aspect of the forearm. Note that this was not necessary for the operation, but provided FPL tendon access for characterization of biological thumb joint biomechanics. **b.** Biological thumb is amputated between the IP and MCP joints (mid-proximal-phalanx). **c.** Intramedullary canal of the proximal phalanx is exposed. **d.** Bone anchor on proximal end of device is inserted into the residual proximal phalanx. **e.** Residual distal end of the FPL tendon is reattached to base of residual proximal phalanx. **f.** Flexion of biological thumb through FPL and FPB (flexor pollicis brevis) is verified. **g.** Reverse radial forearm (RRF) flap is outlined. **h.** RRF flap is harvested and tubularized over the device. **i.** First dorsal metacarpal artery (FDMA) flap is outlined. **j.** FDMA flap is harvested and placed over the biosynthetic thumb tip. **k.** Both flaps are sutured to the native surrounding skin. **l.** Flaps and defects are sutured closed and operation is finished.

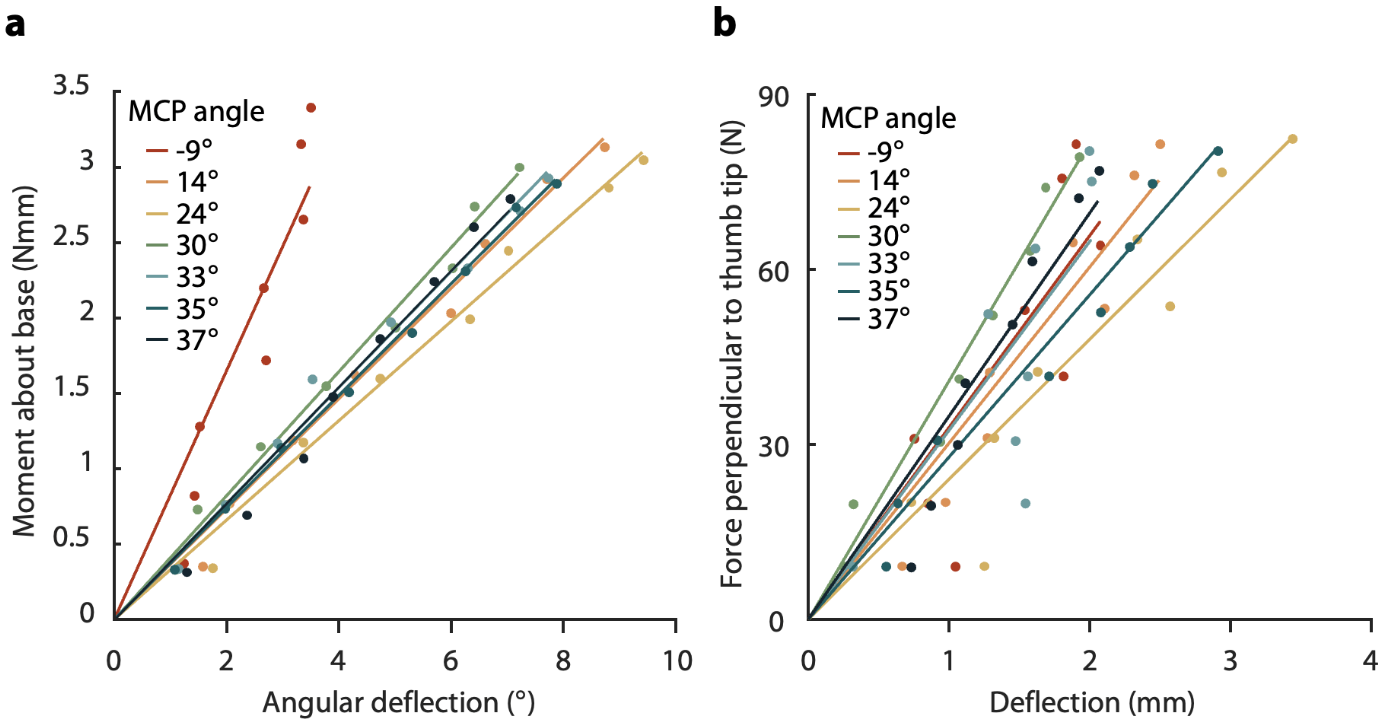


**Supplemental Figure 2.** Thumb-tip deflection under loading. Dots show average measured deflection over a 1.5 second window, beginning after the system reached equilibrium. Lines represent a first-order linear regression. **a.** Moment about the proxy MCP versus angular deflection of the thumb tip. **b.** Force perpendicular to the thumb tip versus thumb tip deflection.

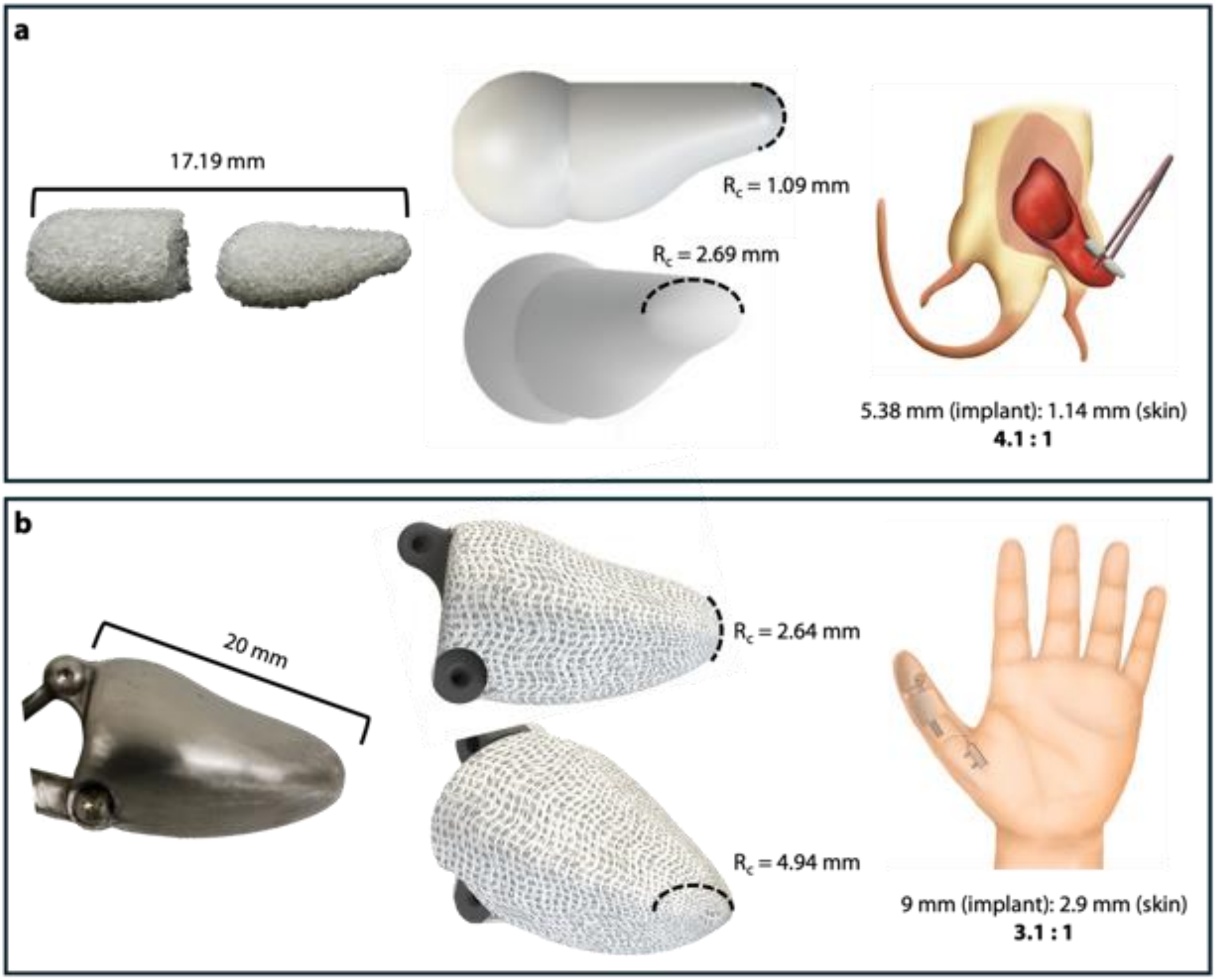


**Supplemental Figure 3.** Two-part implant design for analogous animal model. **a.** Rodent implant and average skin thickness dimensions. **b.** Human implant thumb tip and skin thickness dimensions. The rodent implant was designed to have both a sharper radius of curvature and a larger implant-size-to-skin-thickness ratio than the human model implant, to model a worst-case scenario for extrusion.[59]

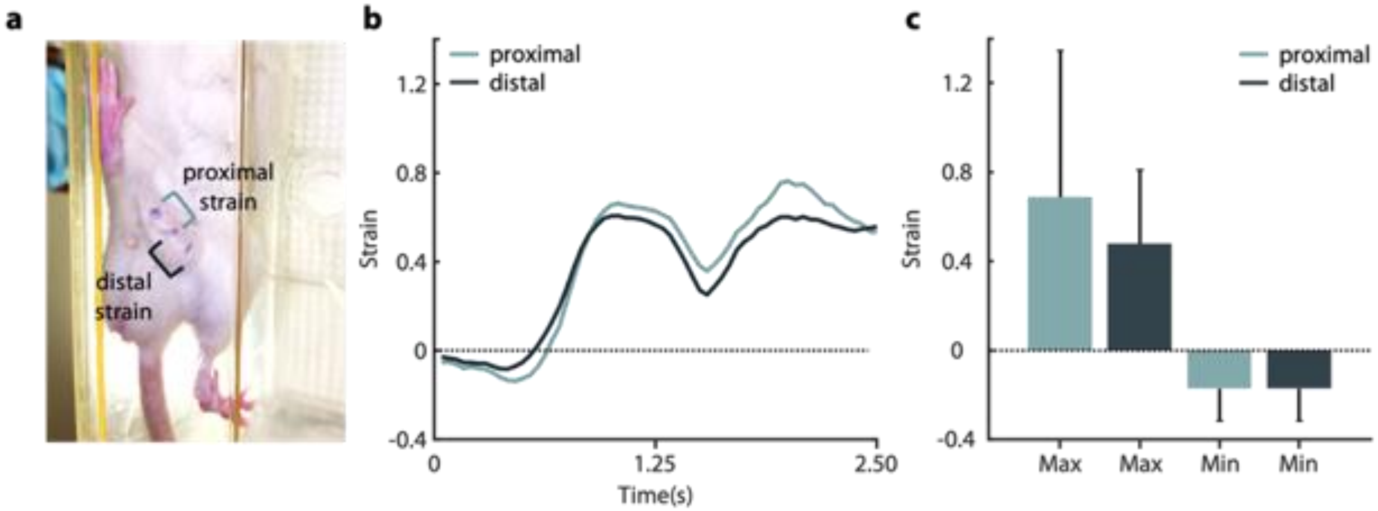


**Supplemental Figure 4.** Rodent flap strain during free walking. **a.** Single frame of a rat subject's epigastric region, taken from the videos used to track flap motion. Three marks were created on the skin flaps: medial and central marks defined the proximal section, and central and distal marks defined the distal section. Video was recorded from beneath as the rats walked across a clear cage. **b.** Proximal and distal strain for a single rat subject (same subject as in part a) versus time. **c.** Average maximum and minimum strain in the proximal and distal regions, across all rat subjects.

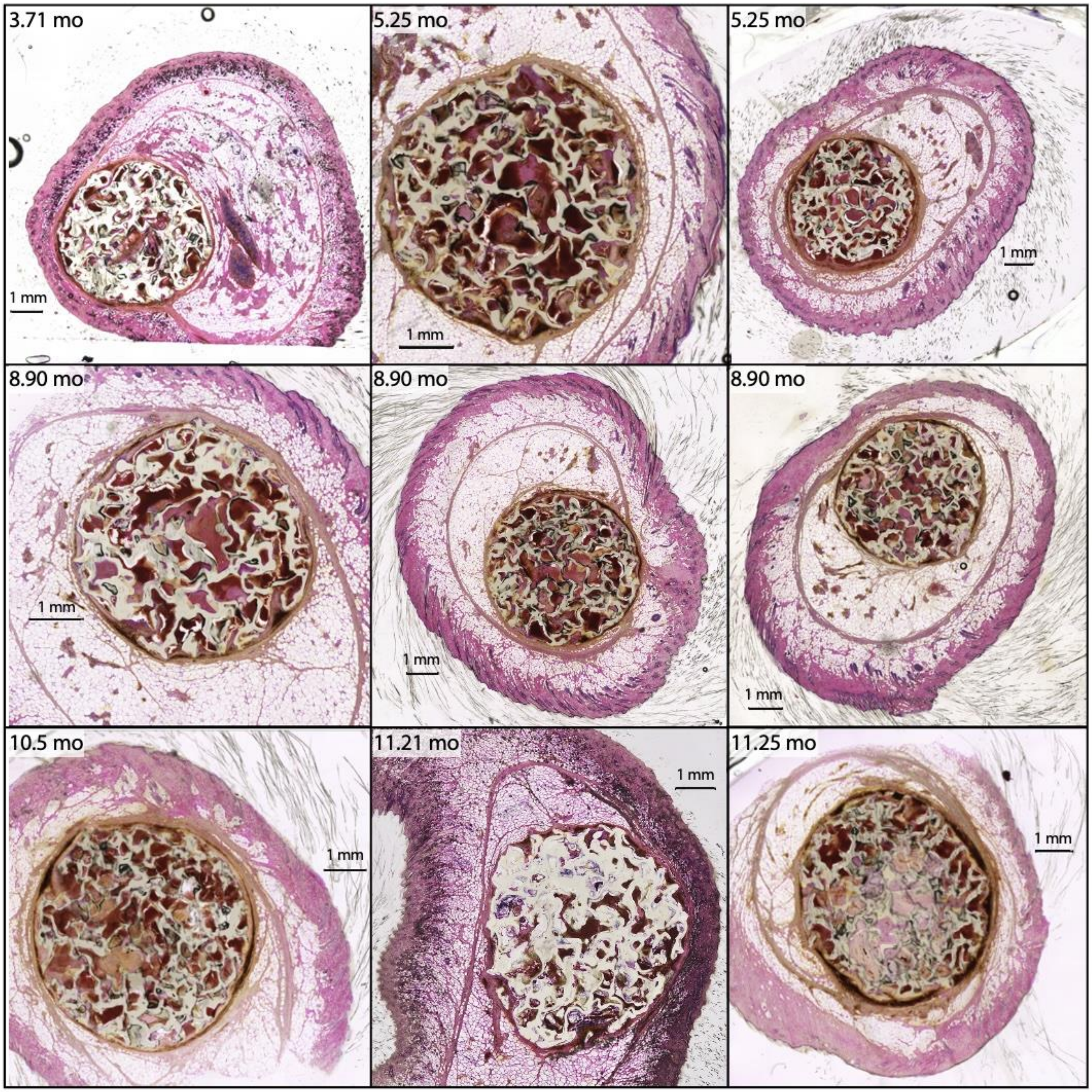


**Supplemental Figure 5.** Histology of the skin flaps from the 9 rat subjects with implants (experimental group). The images are arranged in increasing duration of time that the implant remained in the flap prior to tissue harvesting (left to right, top to bottom).

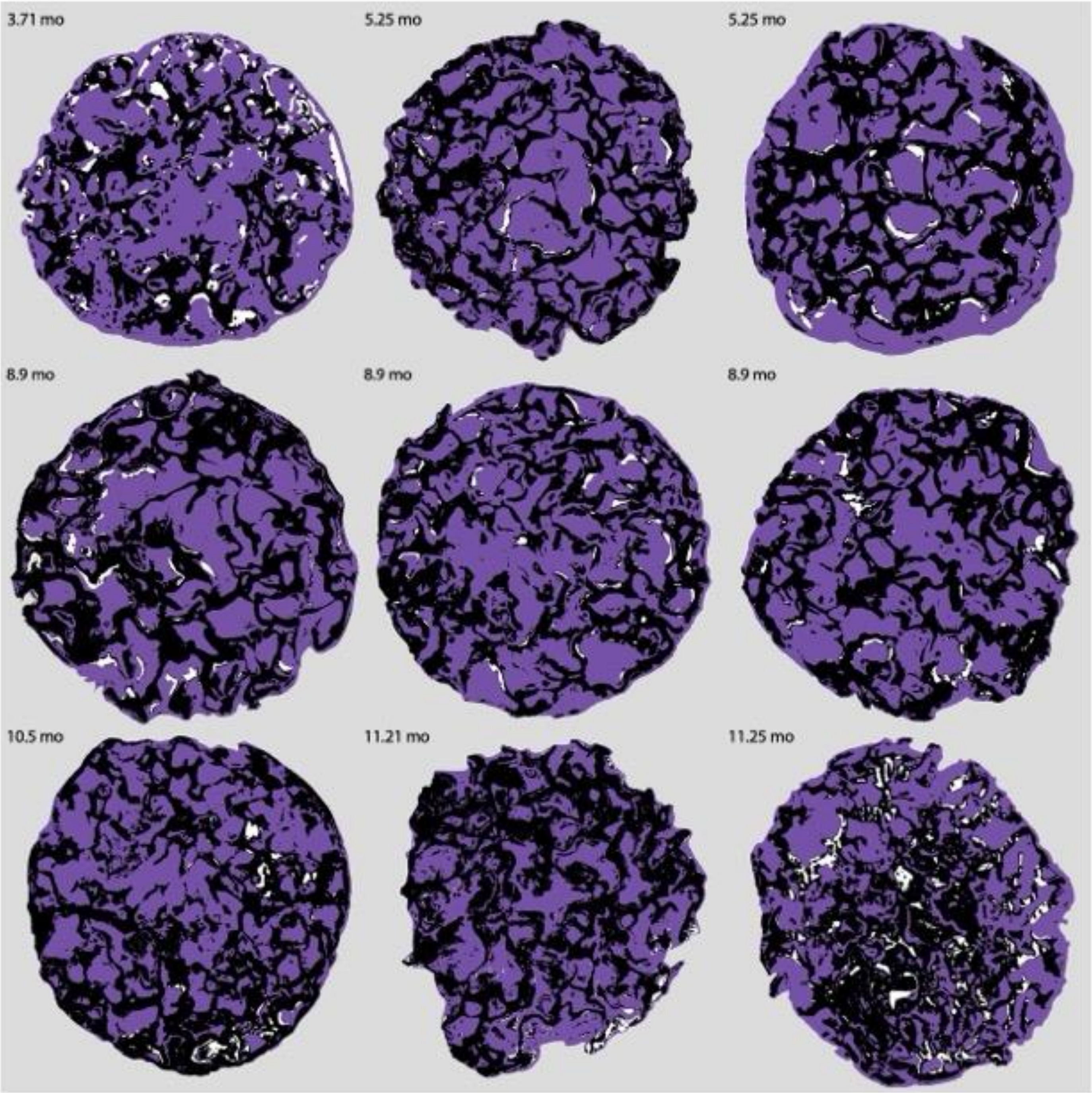


**Supplemental Figure 6.** Histology image processing results. Segmentation is shown, delineating tissue (purple), implant (black), and empty pore space (white) from the histology slides in Supplemental Fig. 5. Images are arranged in the same order as in Supplemental Fig. 5 (harvested from 3.71 through 11.25 months post-surgery).

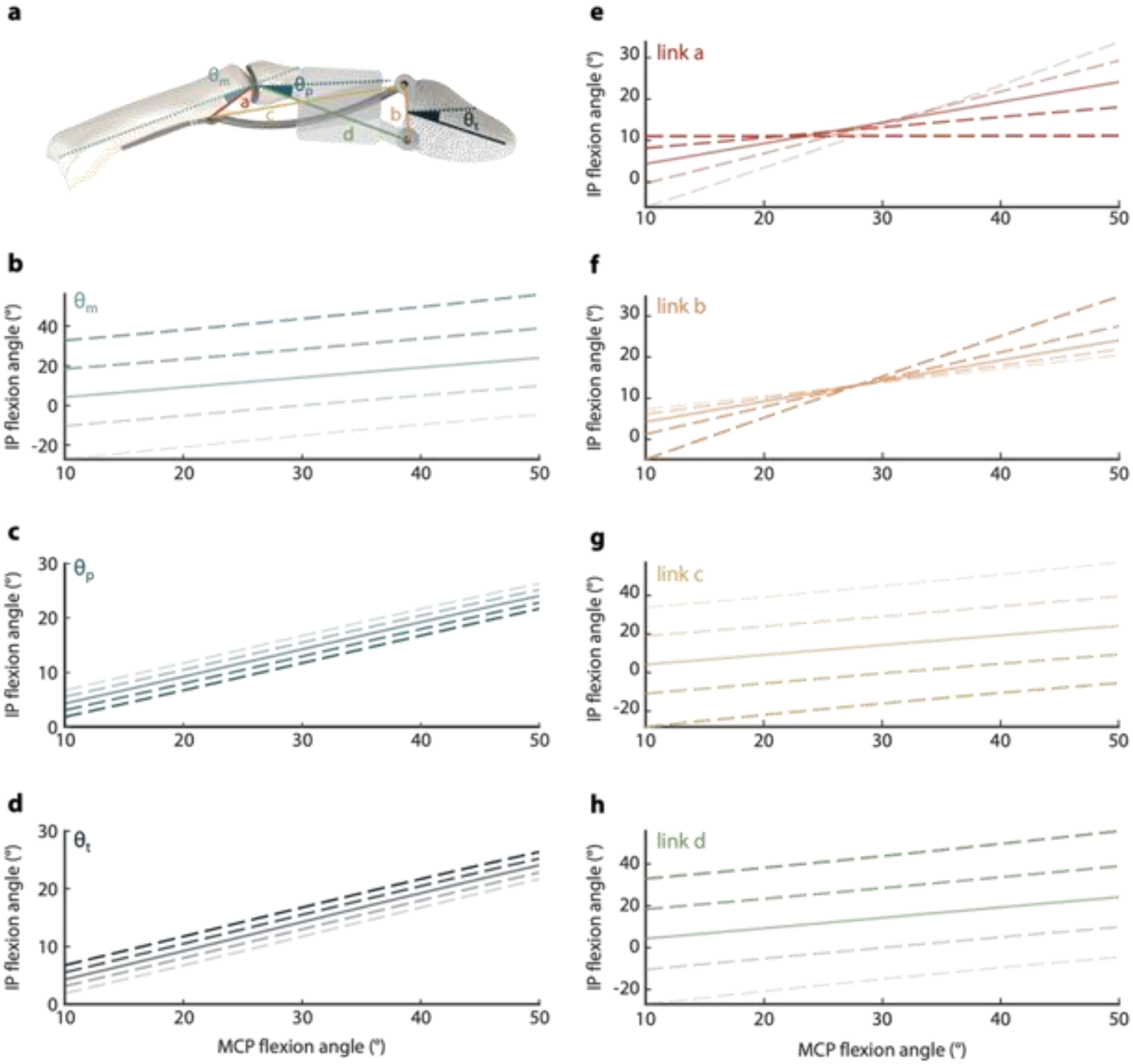


**Supplemental Figure 7.** Sensitivity of crossed four-bar linkage kinematics to design and implantation parameters. **a.** Diagram showing linkage parameters. The linkage is defined by seven parameters that include three angles ($\theta_m$, $\theta_p$, and $\theta_t$) and four link lengths (a, b, c, and d). $\theta_m$ is the angle between link a and the metacarpal long axis of the metacarpal bone. $\theta_p$ is the angle between link d and the long axis of the residual proximal phalanx. $\theta_t$ is the angle formed between a line perpendicular to link b and the thumb tip. **b-d.** MCP-IP angle relationships as each angle parameter is systematically varied around its nominal "clinically viable" value, while all other parameters are held at their nominal values. Nominal performance is shown on each plot by a solid line. Dashed lines represent offsets of -5°, -2.5°, 2.5°, and 5° (dark to light) from the nominal parameter value. **e-h.** MCP-IP angle relationships as each link length parameter is systematically varied around its nominal "clinically viable" value, while all other parameters are held at their nominal values. Nominal performance is shown on each plot by a solid line. Dashed lines represent offsets of -5 mm, -2.5 mm, 2.5 mm, and 5 mm (dark to light) from the nominal parameter value.

| Experiment | Group | Description |
|---|---|---|
| linkage optimization | biological | MCP-IP angle relationship of a biological thumb (ref. (5) main text) |
| | idealized | MCP-IP angle relationship following optimization of mechanical linkage to match "biological" above |
| | clinically viable | MCP-IP angle relationship following constraint to reduce extrusion risk |
| | physical prototype | MCP-IP angle relationship of the manufactured prototype |
| animal study | experimental | rats with flap wrapped around proxy implant |
| | control | rats with flap wrapped without proxy implant |
| cadaver experiment | biological | cadaver with native thumb intact |
| | biosynthetic | cadaver with biosynthetic thumb implant |

**Supplemental Table 1.** Nomenclature used throughout the manuscript. Group names are consistent with those referenced in the main text and figures.
